# Cantu syndrome-associated SUR2 (*ABCC9*) mutations in distinct structural domains result in K_ATP_ channel gain-of-function by differential mechanisms

**DOI:** 10.1101/209783

**Authors:** Conor McClenaghan, Alex Hanson, Monica Sala-Rabanal, Helen I. Roessler, Dragana Josifova, Dorothy K. Grange, Gijs van Haaften, Colin G. Nichols

## Abstract

The complex cardiovascular disorder Cantu Syndrome arises from gain-of-function mutations in either *KCNJ8* or *ABCC9*, the genes encoding the Kir6.1 and SUR2 subunits of ATP-sensitive potassium (K_ATP_) channels. Recent reports indicate that such mutations can increase channel activity by multiple molecular mechanisms. In this study, we determine the mechanism by which K_ATP_ function is altered by several mutations in distinct structural domains of SUR2: D207E in the intracellular L0-linker and Y985S, G989E, M1060I, and R1154Q/W in TMD2. Mutations were engineered at their equivalent position in rat SUR2A (D207E, Y981S, G985E, M1056I and R1150Q/W) and functional effects were investigated using macroscopic rubidium (^86^Rb^+^) efflux assays and patch clamp electrophysiology. The results show that D207E increases K_ATP_ activity by increasing intrinsic stability of the open state, whilst the cluster of Y981S/G985E/M1056I mutations, as well as R1150Q/W, augment Mg-nucleotide activation. The response of mutant channels to inhibition by the sulfonylurea drug glibenclamide, a potential pharmacotherapy for CS, was also tested. There was no major effect on glibenclamide sensitivity for the D207E, Y981S, G985E or M1056I mutations, but glutamine and tryptophan substitution at R1150 resulted in significant decreases in potency.

## Introduction

ATP-sensitive potassium (K_ATP_) channels are potassium-selective ion channels formed by obligate co-assembly of pore-forming Kir6.x subunits and regulatory sulfonylurea receptors (SURx), in a 4:4 stoichiometry [1–4]. Channel opening is dynamically regulated by intracellular nucleotides and membrane phospholipids, and thereby couples the membrane potential of excitable cells to their metabolic state [5]. By binding to, and stabilizing, closed states of the Kir6.x subunit, ATP decreases channel open probability whilst magnesium-nucleotide complexes (MgADP and MgATP) bind to the nucleotide binding domains (NBDs) of SURx subunits to activate the channel [6, 7].

In cardiac, smooth, and skeletal muscle, SUR2 subunits (of which there are two main splice variants, SUR2A and SUR2B) co-assemble variously with Kir6.1 (as in vascular smooth muscle) or Kir6.2 (as in ventricular and skeletal muscle) [8, 9]. In the heart, Kir6.2/SUR2A K_ATP_ channels have been proposed to be critical for ischemic pre-conditioning, whilst in skeletal muscle Kir6.2/SUR2A channels may provide a “brake” to hyperpolarize the membrane potential despite elevations in intracellular calcium during periods of exercise and increased metabolism [9, 10]. In smooth muscle, Kir6.1/SUR2B K_ATP_ activity is a key determinant of electrical excitability and consequent contractility in blood and lymphatic vessels, as well as in bladder and uterine muscle [11–15]

There have now been multiple reports of mutations in the *ABCC9* and *KCNJ8* genes (which encode for SUR2 and Kir6.1 respectively) associated with the complex heritable disorder, Cantu Syndrome (CS) [16–21]. CS patients exhibit diverse cardiovascular features including: dilated and tortuous vessels, cardiomegaly, electrophysiological alterations in the cardiac conduction system, decreased neuro-vascular coupling and persistence of fetal circulation [16–18, 20–25]. An emerging model for the molecular basis of CS is that missense mutations in *ABCC9* or *KCNJ8* result in increased K_ATP_ channel activity, and consequent reduced smooth muscle excitability and contractility [26, 27]. CS-associated mutations in SUR2 have previously been shown to result in K_ATP_ channel gain of function (GoF) by distinct mechanisms, including enhanced Mg^2+^-nucleotide activation and increased intrinsic open probability with consequent decreases in ATP inhibition [18, 19]. Here we examined the functional effects of previously uncharacterized CS mutations that are predicted to cluster together at the link between NBD1 and TMD2: Y981S (human Y985S), G985E (G989E) and M1056I (M1060I) (Fig. 1), and compared the molecular consequences to those of D207E (D207E), located in the intracellular L0-linker, between TMD0 and TMD1 (Fig. 1). In addition, the sensitivity of mutant channels to the sulfonylurea K_ATP_-inhibitor, glibenclamide was tested. Glibenclamide holds promise as a potential treatment for CS, however numerous K_ATP_ GoF mutations which reduce sulfonylurea sensitivity have previously been reported [28–30]. Therefore, determining sulfonylurea sensitivity for specific mutations may be required for future individualized therapy. The results are interpreted alongside structural insights from recently reported high resolution cryo-EM structures of K_ATP_ channel complexes [3, 4] to provide further detail of the molecular basis of K_ATP_ channel GoF in CS.

**Figure 1:**
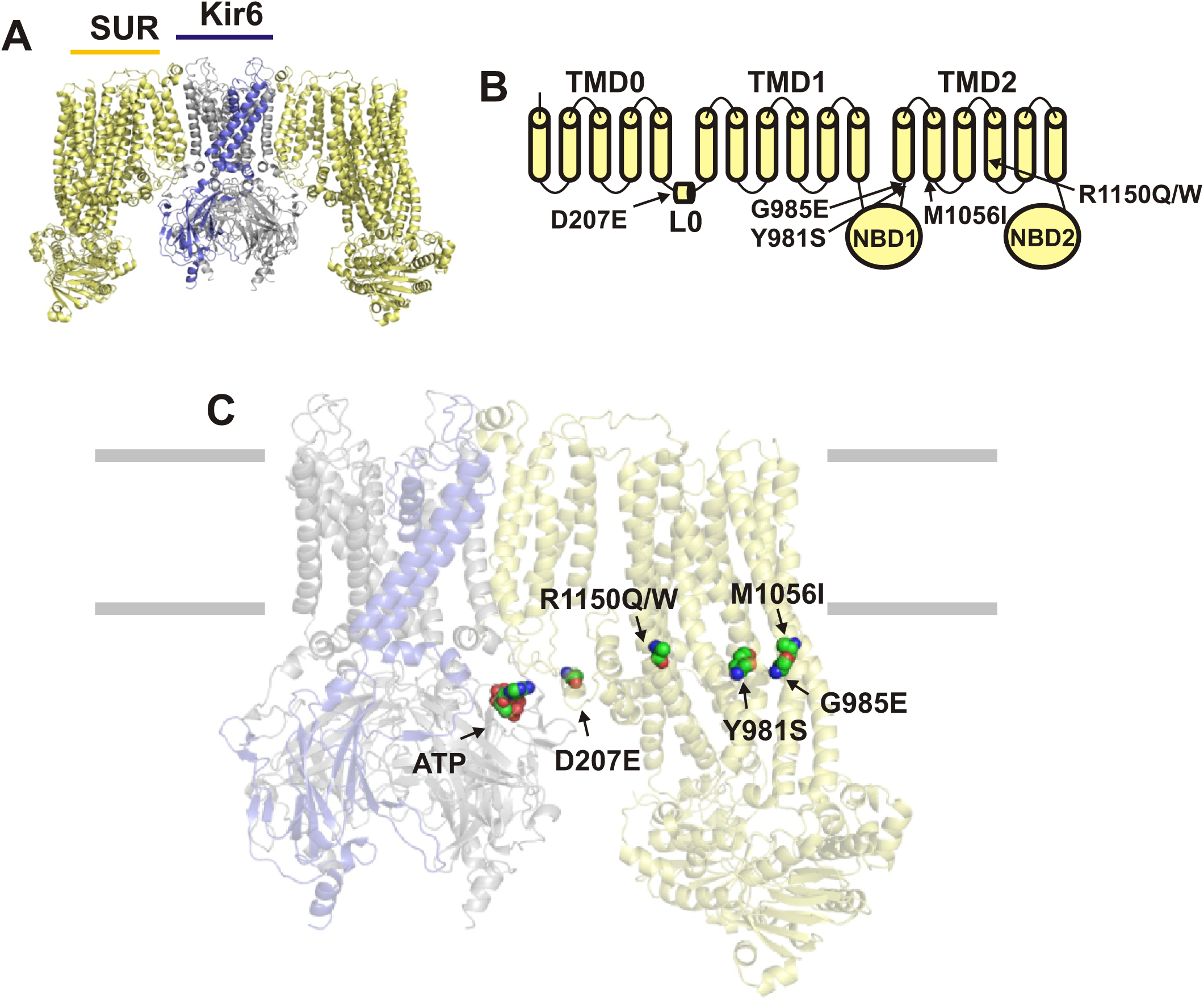
The structure of K_ATP_ channels. **(A)** K_ATP_ channels form as hetero-octamers of four pore-forming Kir6.x subunits each associated with a SUR subunit (two SUR subunits omitted from figure). **(B)** Schematic representation of the position of D207E, Y981S, G985E, M1056I, and R1150Q/W in the linear sequence of SUR2. **(C)** The expected positions of D207E, Y981S, G985E, M1056I, and R1150Q/W mapped onto the pancreatic K_ATP_ (Kir6.2/SUR1) structure [4]. The residues shown are the analogous positions in hamster SUR1 (D209, Y1004, A1008, T1089 and R1183 respectively). ATP is modelled in the Kir6.2 binding site.

## Methods

### Molecular biology and cell culture

Mutations were introduced into a rat SUR2A (pCMV_rSUR2A) cDNA construct using site-directed mutagenesis and verified by direct Sanger sequencing. Cosm6 cells were cultured in Dulbecco’s Modified Eagle’s Medium (DMEM) and transfected using Fugene6 (Roche) with wild-type pcDNA3.1_mKir6.2 (0.6 µg) and wild-type or mutant pCMV_rSUR2A constructs (1 µg) in addition to 0.2 µg of pcDNA3.1_eGFP for visual detection of successful transfection. To model heterozygous expression of mutant subunits, cells were transfected with WT Kir6.2 along with a 1:1 ratio of WT and mutant rSUR2A (0.5 µg: 0.5 µg). Radioactive rubidium efflux experiments were performed 36 h post transfection whilst excised patch clamp recordings were made 48-72 h post transfection.

### ^86^Rb^+^ efflux assay

Transfected Cosm6 cells were plated in 12-well plates to reach 70-80% confluence on the day of experimenting. Prior to commencing the efflux assay the culture medium was replaced by DMEM supplemented with 1 µCi/ml ^86^RbCl (PerkinElmer) and incubated for > 6 hours (37 °C/5% CO_2_) to load the cells with the ^86^Rb^+^ isotope. After the loading incubation, cells were washed with Ringer’s solution containing (in mM) 118 NaCl, 10 HEPES, 25 NaHCO_3_, 4.7 KCl, 1.2 KH_2_PO_4_, 2.5 CaCl_2_, and 1.2 MgSO_4_ either alone or supplemented with 2.5 mg/ml oligomycin and 1 mM 2-deoxy-D-glucose to induce metabolic inhibition (MI) and incubated at room temperature for a further 10 minutes. Cells were then washed three times with Ringer’s solution (either with or without MI supplements) before the experiment was commenced. Ringer’s solution was added to each well, collected, and replaced at the defined time points (2.5, 5, 12.5 and 22.5 and 37.5 minutes). After the experiment, cells were lysed with 2% SDS to attain the remaining intracellular ^86^Rb^+^ and sample radioactivity was determined by scintillation counting.

The cumulative ^86^Rb^+^ efflux at each time point was calculated from the total counts from each sample (including the ^86^Rb^+^ remaining post-cell lysis). Apparent K_ATP_-independent efflux rate constants (*k*_*1*_) were obtained from GFP-transfected cells using the equation:

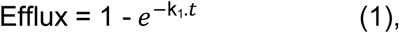

and K_ATP_-dependent efflux rate constant (*k_2_*) was obtained from K_ATP_-transfected cells using the equation:

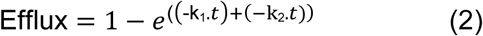

where k_1_ was obtained from GFP-transfected cells (equation 1). The number of active channels was assumed to be proportional to *k_2_.* In MI conditions a time-dependent divergence from a mono-exponential efflux is observed. This is attributed to inactivation of background efflux mechanisms over time, therefore in this condition rate constants were derived from exponential functions fit to early time points only (2.5 – 12.5 min). Data shown represents the mean ± S.E.M. from at least 3 independent experiments each with multiple replicates (N ≥ 3, n ≥ 7). Statistical significance was determined using Mann-Whitney U tests with a p value < 0.05 deemed statistically significant.

### Inside-out excised patch clamp recordings

Pipettes were made from soda lime glass microhematocrit tubes (Kimble) and had a resistance of 1 – 2 MΩ when filled with pipette solution.The bath and pipette solutions (K_INT_) contained (in mM): 140 KCl, 10 HEPES, 1 EGTA (pH 7.4 with KOH). Currents were recorded at a constant holding potential of −50 mV in the absence and presence of nucleotides as indicated. Where included, free Mg^2+^ concentrations were maintained at 0.5 mM by supplementation of MgCl_2_ as calculated using CaBuf (Katholieke Universiteit Leuven).

Where stated, porcine brain L-α-phosphatidylinositol-4,5-bisphosphate (PIP_2_) (Avanti Polar Lipids) was applied at 5 µg/ml. Rapid solution exchange was attained using a Dynaflow Resolve perfusion chip (Cellectricon). Experiments were performed at 20-22 °C. K_ATP_ channel currents in solutions of varying nucleotide concentrations were normalized to the basal current in the absence of nucleotides and dose-response data were fit with a four-parameter Hill fit according to the equation below:

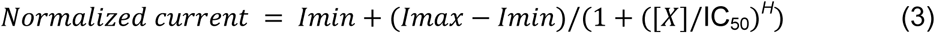

Where the current in K_int_ = *I*_*max*_ = 1, *I*_*min*_ is the normalized minimum current observed in ATP/MgATP/glibenclamide, [*X*] refers to the concentration of ATP/MgATP/glibenclamide, *IC*_*50*_ is the concentration of half-maximal inhibition and *H* denotes the Hill coefficient.

Data were tested for statistical significance using the Mann Whitney U test, and presented as mean ± S.E.M.

## Results

### Case history of subject with SUR2[Y985S] mutation

The subject is the fourth child of healthy, unrelated Caucasian parents, with no family history of relevance to her condition. The pregnancy was complicated with raised nuchal translucency at twelve weeks and polyhydramnios at thirty-two weeks gestation. At thirty-eight weeks of gestation, labour was induced, with uncomplicated vaginal delivery. Birth weight was 5.3kg (>99 centile). There was no significant delay in early development, but language skills developed slowly. At birth, hypertrichosis was evident, with a full head of dark hair with low anterior hairline, shoulders, arms, legs and back were covered with long, thick and dark hair. At three years of age, facial features were rather coarse, with mild epicanthic folds and down slanting palpable fissures with full lips and a broad face. The forehead was extremely low with fine hair in front of the ears, extending over her chin and lanugo over her neck and chest. The heart was slightly enlarged, but there was no overt evidence of cardiomyopathy.

At age 5, the subject presented with recurrent respiratory infections and required hospital admission for pneumonia, leading to tonsillectomy and adenoidectomy, which improved severe snoring and obstructive sleep apnoea. Height was on the 50th centile, weight on the 91st centile and her head circumference was on the 98th centile. Facial features remained coarse with down slanting palpable fissures, full cheeks, broad tip to the nose with mild thickening of the alae nasae and a low columella. Significant joint laxity was evident in the hands, with deep palmar creases and soft skin on the palms and generous fetal finger pads.

This subject thus exhibited most of the features typically found in individuals with CS [18, 24]. Sequencing of *ABCC9* coding regions revealed a heterozygous mutation (c.2954A>C, p.Y985S) that was absent in genomic DNA from either parent. Heterozygous *de novo* mutations (p.G989E; p.M1060I) were also identified in two additional diagnosed CS subjects, for whom clinical details are not available. The three mutated residues are predicted to cluster in a similar location within the SUR2 protein (Fig. 1). In the following study we therefore analyzed the molecular consequences of these mutations, and compared them to the consequences of the most common CS mutation (p.R1154Q), and another uncharacterized CS mutation p.D207E [16], located in distinct SUR2 domains.

### Cantu Syndrome mutations result in gain of function of K_ATP_ channel in intact cells

To determine the effect of mutations on K_ATP_ channel function, SUR2A constructs were co-expressed with Kir6.2 in Cosm6 cells, and channel activity assessed using a radioactive 86Rb^+^ flux assay. Firstly, basal K_ATP_ activity under quasi-physiological regulation by intracellular nucleotides in intact cells was determined by measuring ^86^Rb^+^ efflux from cells bathed in Ringer’s solution. As shown in Figs. 2A and B, the K_ATP_-dependent ^86^Rb^+^ efflux rate in these conditions was significantly increased by each mutation, confirming the expected GoF effect. Efflux rates in basal conditions are a function of both channel activity and surface membrane expression. We assessed efflux rates in cells subjected to metabolic inhibition (MI) to decrease intracellular ATP-synthesis and relieve channels of nucleotide regulation. As shown in Figs. 2C and D, there was a statistically insignificant decrease in Y981S efflux rate, and a statistically insignificant increase in MI efflux rate was observed for D207E, which may indicate that these mutations result in a small decrease or increase, respectively, in plasma membrane expression in this heterologous system. Otherwise, the maximum flux under these conditions was not markedly different between constructs, suggesting that the number of active channels was similar.

**Figure 2:**
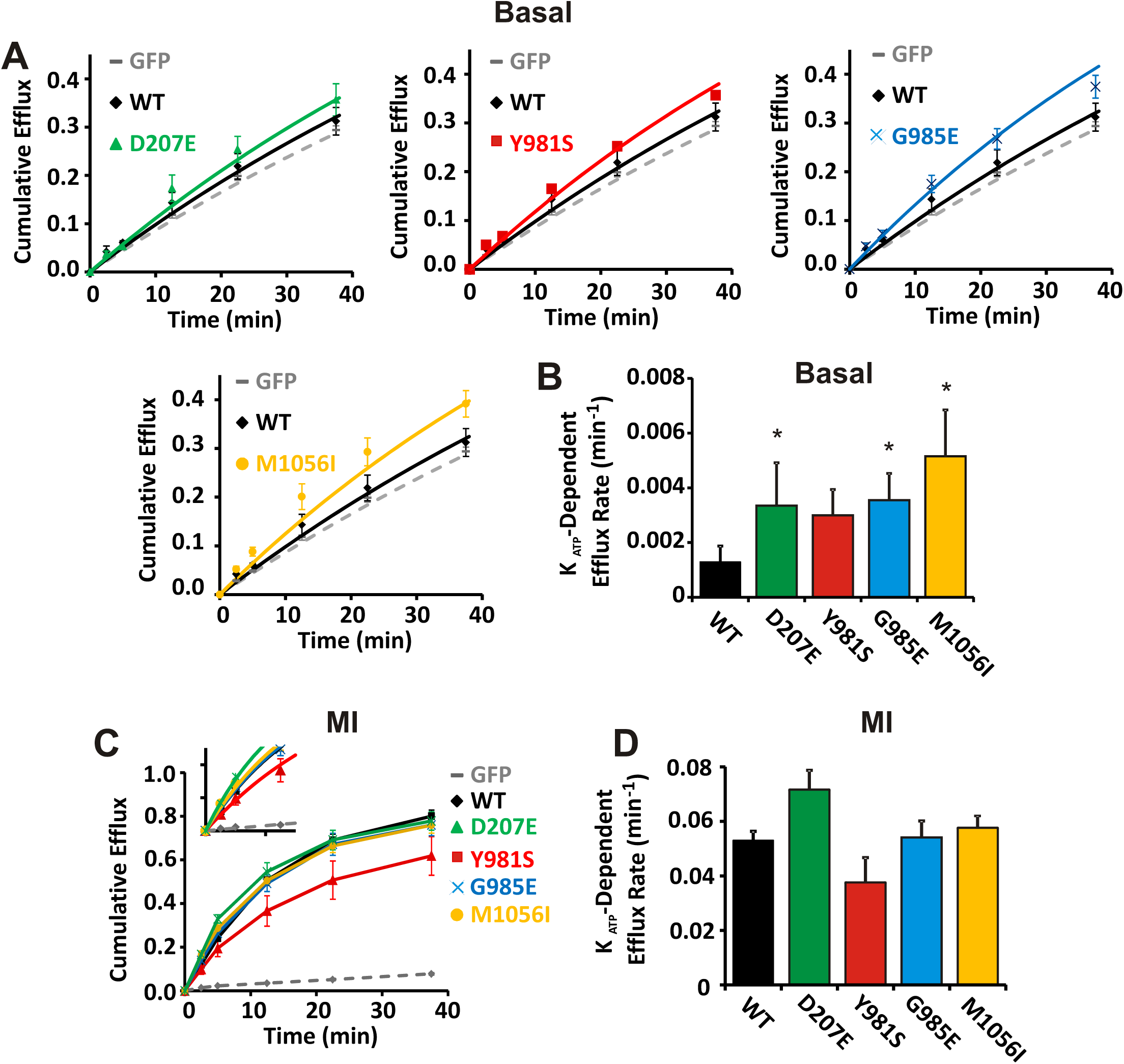
D207E, Y981S, G985E and M1056I increase KATP channel activity. **(A)** Cumulative ^86^Rb^+^ efflux was measured from Cosm6 cells transfected either with GFP alone or with Kir6.2 plus WT or mutant SUR2A. Efflux as a function of time was first recorded in basal conditions (cells incubated in Ringer’s solution) **(A)** and the K_ATP_-dependent efflux rate was attained from exponential fits to efflux time data **(B)**. Secondly, efflux was measured from cells subjected to metabolic inhibition (MI), induced by incubation in Ringer’s solution with oligomycin and 2-deoxy-d-glucose **(C)**, and the rate constant for KATP-dependent efflux in MI was calculated from exponential fits to the early time points **(inset C & D)**. Data and error bars show represents mean ± S.E.M. from 3–5 independent experiments. Statistical significance denoted by asterisk (*) and defined as p < 0.05 according to Mann Whitney U test.

All known CS patients are heterozygous and we modeled heterozygous conditions by co-expressing Kir6.2 together with WT SUR2A and mutant SUR2A subunits at a 1:1 ratio. The resultant channels were assayed by monitoring ^86^Rb^+^ efflux. Only very minor increases in basal efflux rate were observed for D207E, G985E and M1056I, whilst a moderate, statistically significant increase was observed for Y981S channels (Fig. 3). Taken together, these data demonstrate that whilst all tested mutations result in K_ATP_ gain-of-function, in heterozygous conditions the effect is subtle under basal conditions.

**Figure 3:**
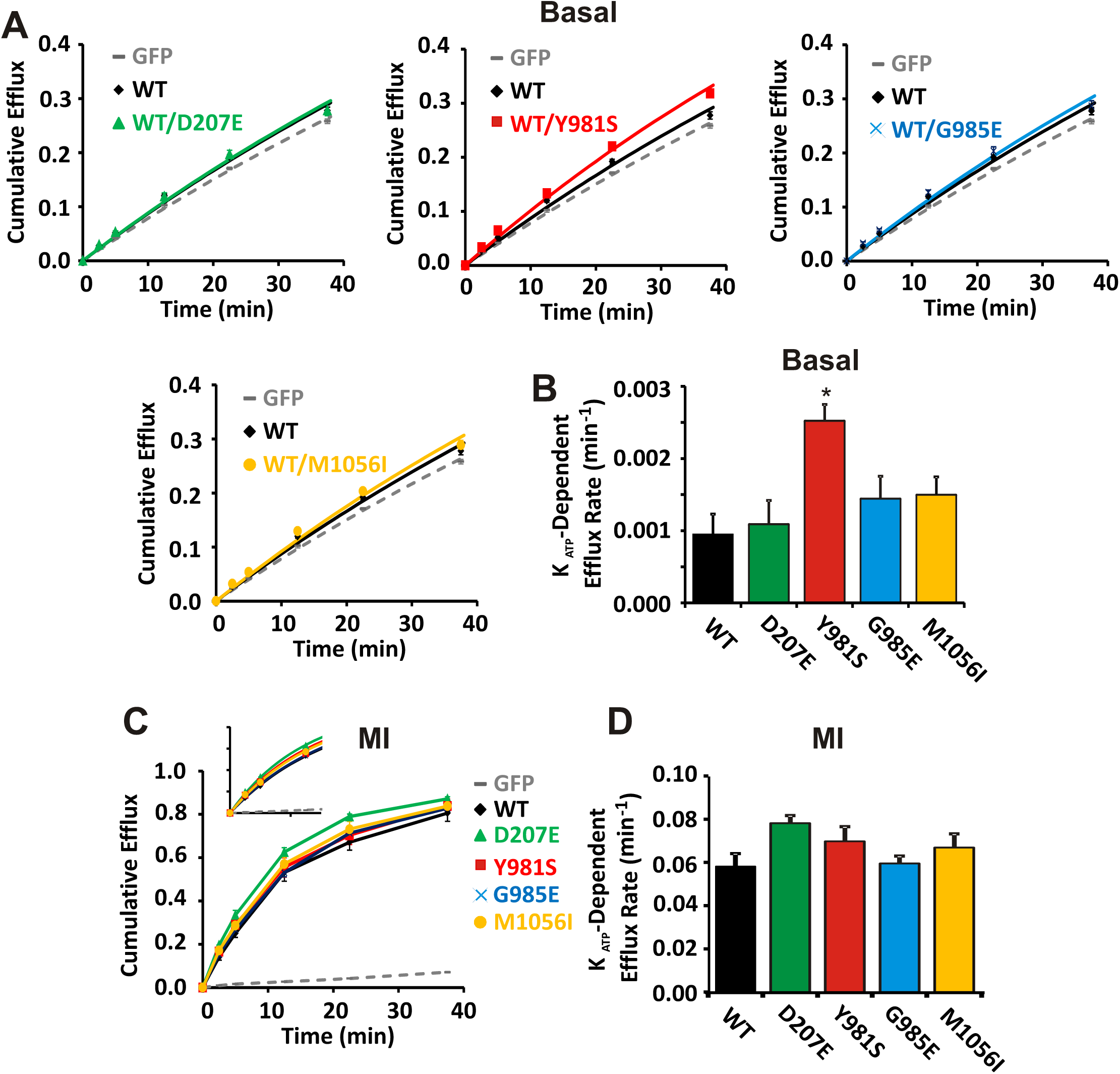
Minor gain-of-function effects of CS-mutant SUR2A when co-expressed with WT subunits in intact cells. Cumulative ^86^Rb^+^ efflux was measured from Cosm6 cells transfected either with GFP alone or with Kir6.2 alongside SUR2A-WT alone, or SUR2A-WT with mutant SUR2A at a 1:1 ratio. Basal efflux shown as function of time **(A)** and K_ATP_-dependent efflux rate constants were calculated **(B)**. Efflux rate was measured in MI conditions **(C)** and rate constants calculated from early time points **(inset C & D).** Data shown represents mean ± SEM from 3 – 5 independent experiments. Statistical significance denoted by asterisk (*) and defined as p < 0.05 according to Mann Whitney U test.

### D207E in the L0 linker increases open-state stability and decreases ATP-inhibition

Increased basal K_ATP_ channel activity could arise from changes in multiple distinct nucleotide sensing mechanisms, including increased Mg-nucleotide activation, or decreased ATP inhibition, either as a result of decreased binding affinity or decreased efficacy due to increase in intrinsic open state stability [19]. To investigate the effects of the above mutations on nucleotide sensitivity, we used inside-out patch clamp recordings. As shown in Fig. 4, channels comprising Kir6.2 and WT SUR2A were inhibited by ATP in the absence of Mg^2+^ with an IC_50_ of ~15 µM. In the presence of Mg^2+^ the IC_50_ was ~20 µM. The presence of Mg^2+^ results in increased channel activity for a given ATP concentration, reflecting mixed effects of the MgATP activation and Mg-independent ATP inhibition, and is reflected in a right-shift in the MgATP dose response curve, relative to Mg^2+^ free ATP. The sensitivity of D207E-containing channels to both MgATP and Mg-free ATP was reduced ~3-fold in each case, with IC_50_ in Mg-free ATP of ~40 µM and ~55 µM, respectively (Fig. 4).

**Figure 4:**
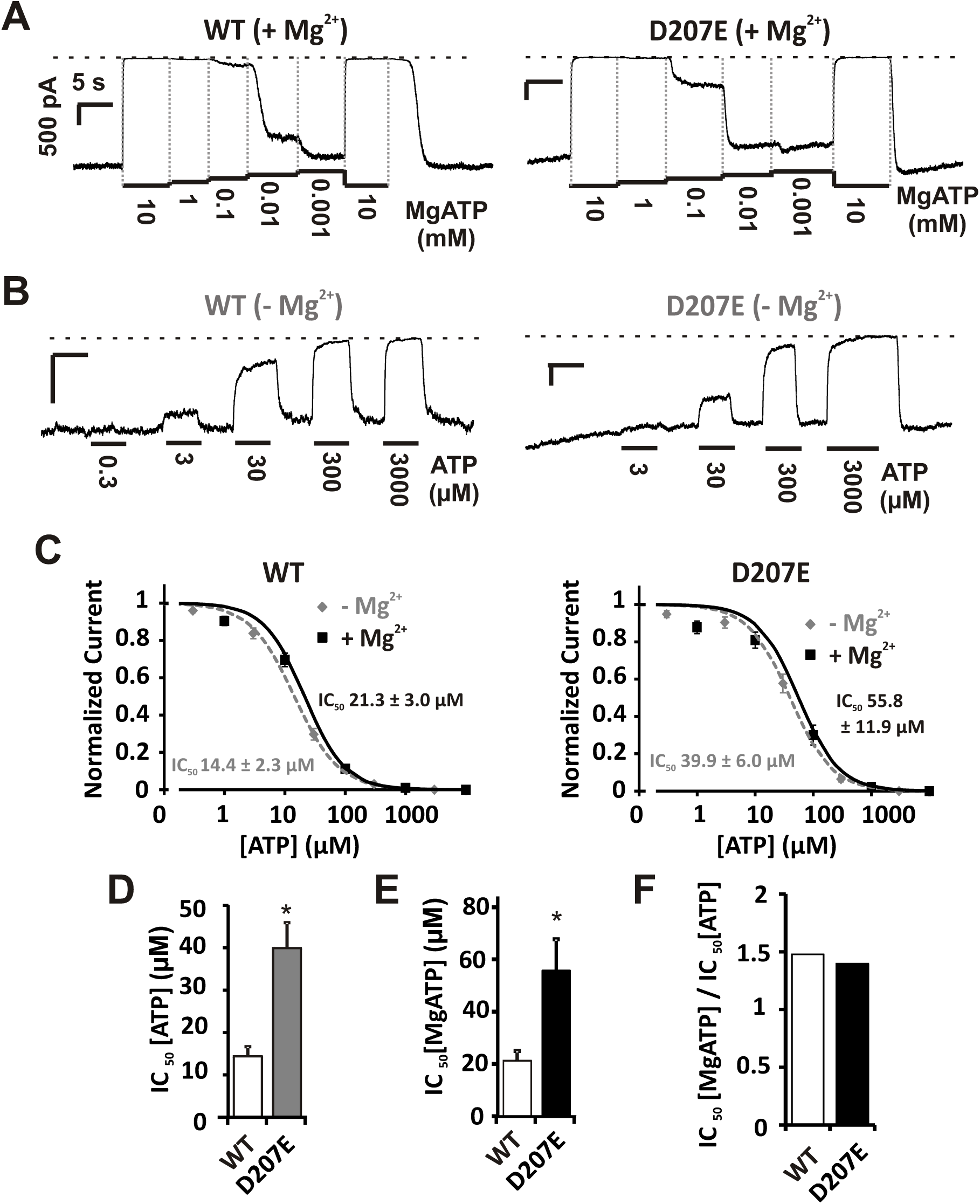
D207E provokes decreases in ATP inhibition in the presence and absence of Mg^2+^. Inside-out patch clamp recordings were made from Cosm6 cells transfected with Kir6.2 plus either SUR2A-WT or D207E. The response to MgATP was determined from voltage clamped patches (−50 mV), as shown in representative traces (scale bars denote 500 pA/5 s unless otherwise stated) **(A)**. ATP inhibition in the absence of Mg^2+^ was also determined from voltage clamped patches (-50 mV), as shown in representative traces **(B)**. Kir6.2/SUR2A-WT channels were inhibited by ATP in the presence of Mg^2+^ with an IC_50_ of 21.3 ± 3.0 µM (Hill coefficient 1.3 ± 0.1; n = 16), and by ATP in the absence of Mg^2+^ with an IC_50_ of 14.4 ± 2.3 µM (Hill coefficient 1.2 ± 0.1; n = 14). This was increased in both cases by the D207E mutation, here, the IC_50_ for ATP in the presence of Mg^2+^ was 55.8 ± 11.9 µM (Hill coefficient 1.2 ± 0.1; n = 9), and by ATP in the absence of Mg^2+^ with an IC_50_ of 39.9 ± 6.0 µM (Hill coefficient 1.2 ± 0.1; n = 9) **(C-E)**. Notably, the ratio of the IC_50_ for ATP in the absence and presence of Mg^2+^ was unchanged by the D207E mutation, indicating the mutation had little effect on Mg^2+^-nucleotide activation **(F)**. Statistical significance denoted by asterisk (*) and defined as p < 0.05 according to Mann Whitney U test.

Thus the relationship between the IC_50_ in ATP in the presence and absence of Mg^2+^ (IC_50_[MgATP]/IC_50_[ATP]) can provide an indirect measure of the extent of Mg^2+^-nucleotide activation, For D207E and WT SUR2A the IC_50_[MgATP]/IC_50_[ATP] was very similar (~1.5), indicating that sensitivity to Mg^2+^-activation was unaffected by this mutation (Fig. 4F), and suggests that GoF results from decreased sensitivity to inhibitory ATP itself. Considering the location of this residue, predicted to be in close proximity to the ATP binding site on Kir6.1 in the octameric K_ATP_ complex [3, 4], this could conceivably arise from altered binding affinity. Alternatively, decreased ATP sensitivity could be the result of enhanced open-state stability of channels [19, 31]. To test the latter directly, we measured the response of channels to PIP_2_ perfused onto the intracellular surface of excised membrane patches (Fig. 5). PIP_2_ increases open probability (Po) to ~1, and the ratio of initial current levels to the activated level in PIP_2_ provides an estimate of the initial ‘intrinsic’ Po [28]. As shown in Fig. 5, the intrinsic Po was increased from ~0.4 in WT to ~0.7 in D207E channels. Therefore, the D207E mutation within the L0 linker results in K_ATP_ gain of function by increasing the intrinsic open probability of channels, rather than by decreasing inhibitory ATP binding affinity.

**Figure 5:**
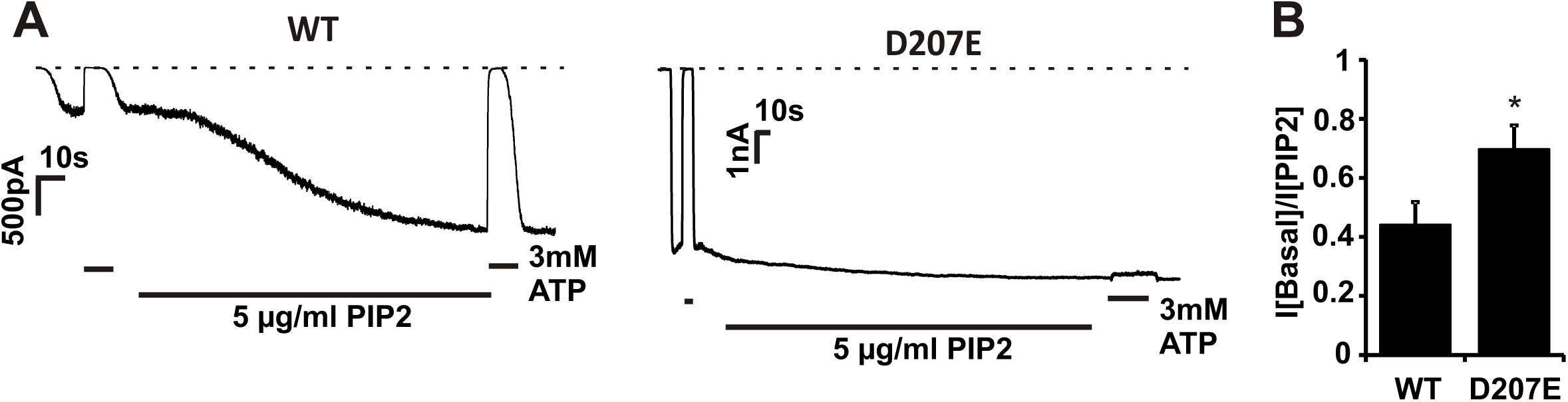
D207E increases intrinsic open-state stability. Inside out patch clamp recordings were made from Cosm6 cells transfected with Kir6.2 alongside either SUR2A-WT or D207E. Patches were administered 3 mM ATP followed by washout and administration of 5 µg/ml PIP_2_ to maximally increase channel activity, as indicated on representative traces **(A)**. The stable peak current observed in PIP_2_ was divided by the current observed In the absence of nucleotides/PIP_2_ to determine the level of basal channel activity. D207E channels displayed increased basal activity (I[Basal]/I[PIP_2_] was 0.44 ± 0.08 for SUR2A-WT and 0.73 ± 0.06 for D207E; data and error bars show mean ± S.E.M. from 7 patches) **(B)**, statistical significance denoted by asterisk (*) and defined as p < 0.05 according to Mann Whitney U test.

### Mutations within the Y981/G985/M1056 cluster increase Mg^2+^-nucleotide activation

The disease-associated mutations Y981S, G985E and M1056I are all clustered togetheron transmembrane helices 12 and 13 in TMD2 (Fig. 1). In comparison to WT SUR2A the IC_50_ for ATP inhibition in the presence of Mg^2+^ was significantly increased by each of these mutations although, in contrast to D207E, there was no effect on ATP sensitivity in the absence of Mg^2+^ (Fig. 6). This is further demonstrated by the increase in IC_50_[MgATP]/IC_50_[ATP] for all mutants (Fig. 6G), indicating that the mutations in this cluster of residues linking NBD1 to TMD2 increase channel activity by enhancing Mg-nucleotide activation.

**Figure 6:**
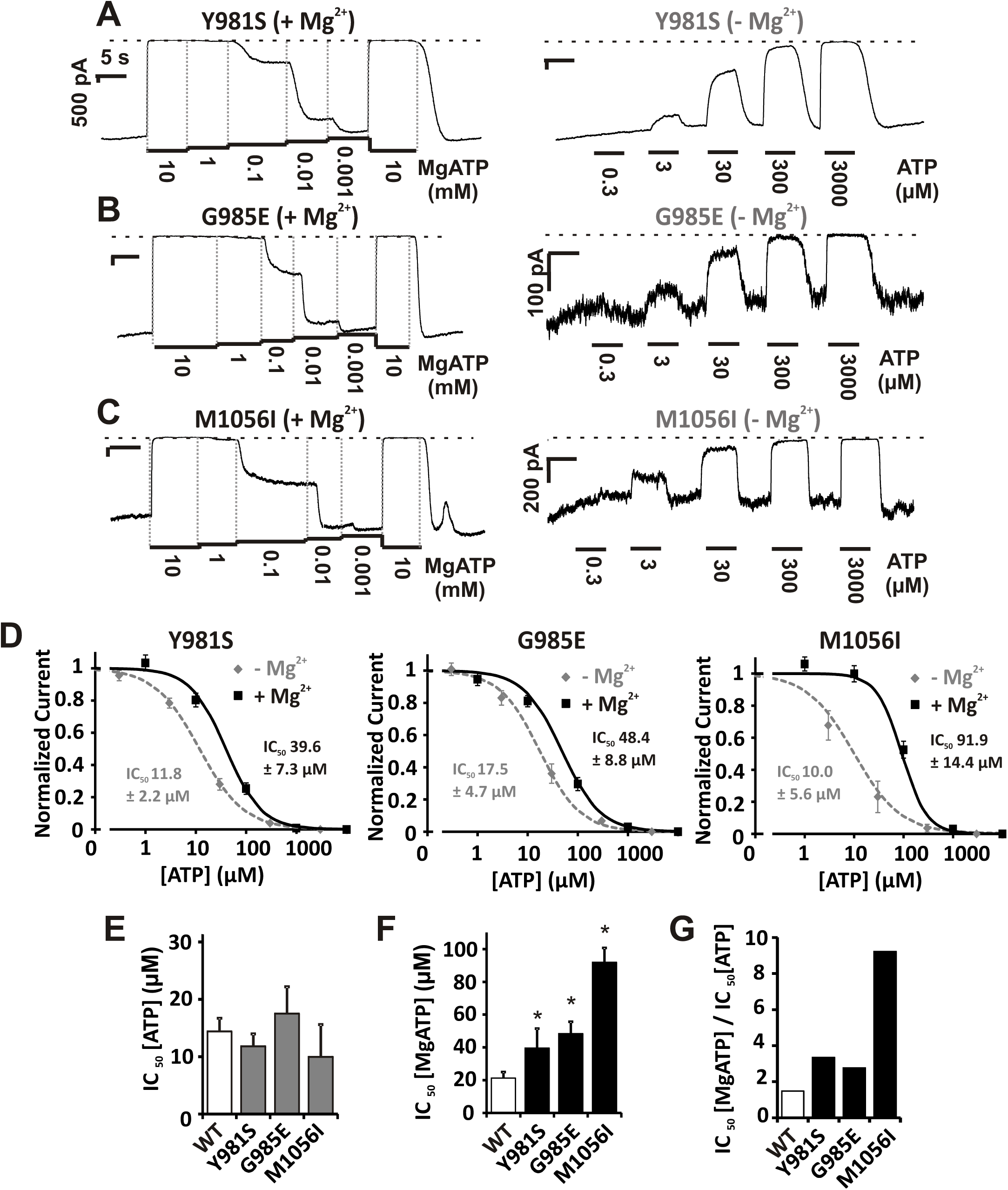
Y981S, G985E and M1056I cause K_ATP_ GoF by enhancing Mg^2+^-nucleotide activation. Inside-out patch clamp recordings were made from Cosm6 cells transfected with Kir6.2 alongside mutant SUR2A. The response to ATP in the presence (left) and absence (right) of Mg^2+^ was determined from voltage clamped patches (-50 mV) of cells expressing either Y981S **(A)**, G985E **(B)** or M1056I **(C)**, as shown in representative traces (scale bars denote 500 pA/5 s unless otherwise stated). Analysis of dose response experiments showed that each mutation increased the IC_50_ for MgATP compared to WT (MgATP IC_50_ for Y981S was 39.6 ± 7.3 µM, Hill coefficient 1.2 ± 0.1, n = 6; MgATP IC_50_ for G985E was 48.4 ± 8.7 µM, Hill coefficient 1.1 ± 0.1, n = 11; MgATP IC_50_ for M1056I was 91.9 ± 14.4 µM, Hill coefficient 1.7 ± 0.2, n = 6), with little effect on ATP inhibition in the absence of Mg^2+^ (ATP IC_50_ for Y981S was 11.8 ± 2.2 µM, Hill coefficient 1.0 ± 0.1, n = 4; ATP IC_50_ for G985E was 17.5 ± 4.7 µM, Hill coefficient 1.1 ± 0.1, n = 6; ATP IC_50_ for M1056I was 10.0 ± 5.6 µM, Hill coefficient 0.9 ± 0.1, n = 3) **(D-F)**. Decreased ATP inhibition in the presence of Mg^2+^ only is demonstrated by the ratio of the IC_50_ for ATP in the presence and absence of Mg^2+^, which is markedly increased for Y981S, G985E and M1056I **(G)**. Statistical significance denoted by asterisk (*) and defined as p < 0.05 according to Mann Whitney U test.

### R1150Q and R1150W in TMD2 also enhance Mg^2+^-nucleotide activation

Having established that the TMD2 Y981S, G985E and M1056I mutations enhance Mg^2+^-nucleotide activation we sought to test whether this mechanism was conserved for other TMD2 mutations, the most common CS-associated mutations R1150Q and R1150W. In agreement with a previous report [16], we show that R1150Q causes a large increase in MgATP IC_50_ whilst R1150W has a more modest effect (Fig. 7). In contrast, R1150Q and R1150W caused only slight increases in ATP IC_50_ (Fig. 7), again reflected by increased IC_50_[MgATP]/IC_50_[ATP] for R1150Q and R1150W (Fig. 7F), and thus both R1150Q and R1150W cause gain of function predominantly by enhancing Mg^2+^-nucleotide sensitivity.

**Figure 7:**
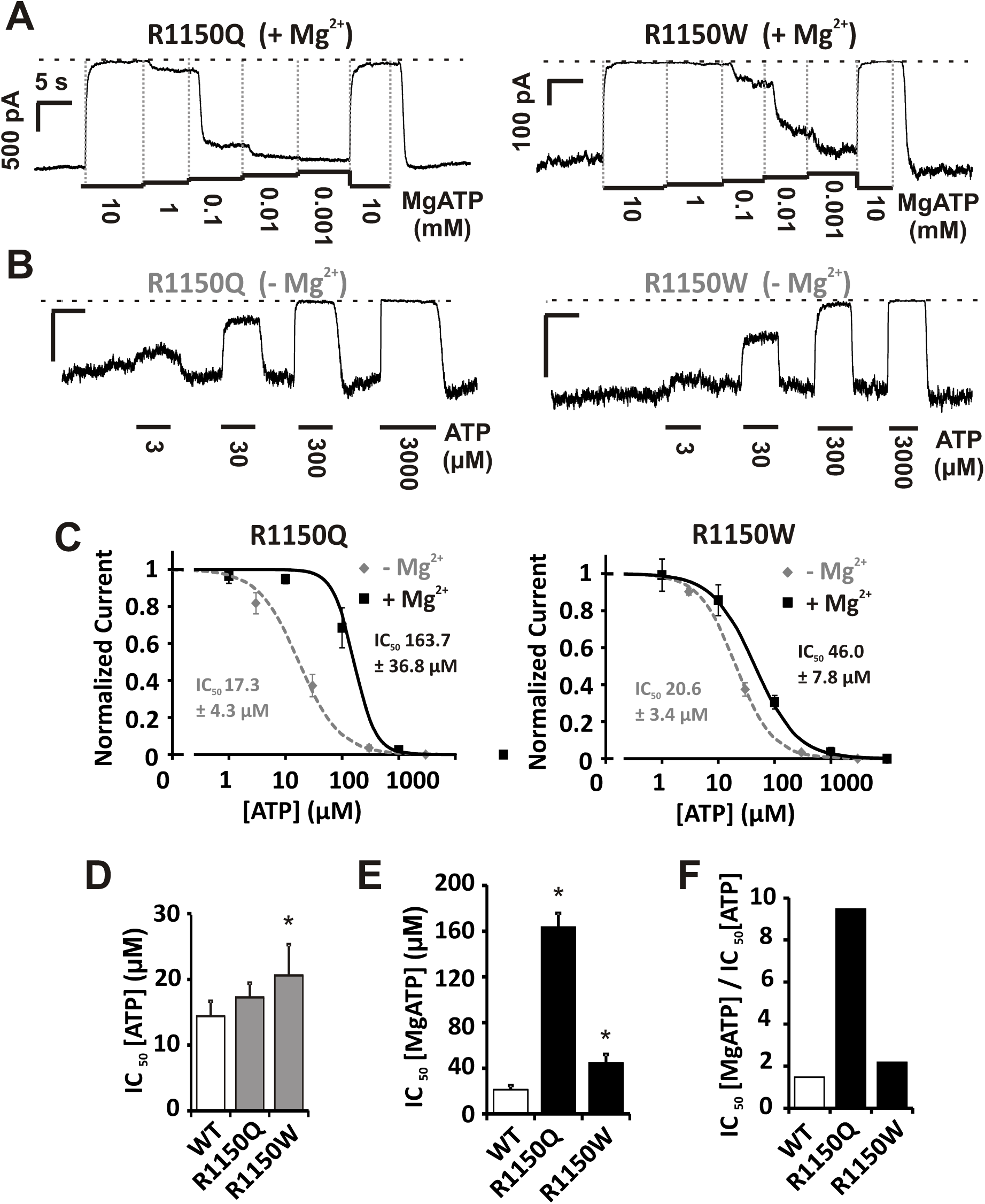
R1150Q and R1150W also cause K_ATP_ GoF by enhancing Mg^2+^-nucleotide activation. The response to MgATP **(A)** and ATP in the absence of Mg^2+^ **(B)** was determined for channels expressing Kir6.2 with R1150Q or R1150W, by inside-out patch clamp recordings from Cosm6 cells as shown in representative traces (scale bars denote 500 pA/5 s unless otherwise stated). Dose response analysis **(C-E)** demonstrated that both mutations markedly increased the IC_50_ for ATP in the presence of Mg^2+^, compared to WT (MgATP IC_50_ for R1150Q was 163.7 ± 36.8 µM, Hill coefficient 2.3 ± 0.3, n = 6; MgATP IC_50_ for R1150W was 46.0 ± 7.8 µM, Hill coefficient 1.2 ± 0.2, n = 6). The effect on ATP sensitivity in the absence of Mg^2+^ was more modest for both mutations (ATP IC_50_ for R1150Q was 17.3 ± 4.3 µM, Hill coefficient 1.2 ± 0.2, n = 8; ATP IC_50_ for R1150W was 20.6 ± 3.4 µM, Hill coefficient 1.3 ± 0.2, n = 10). An increase in the ratio of the IC_50_ for ATP in the presence over the IC_50_ for ATP in the absence of Mg^2+^ indicates that these mutations also predominantly confer GoF by augmentation of Mg^2+^-nucleotide activation. Statistical significance denoted by asterisk (*) and defined as p < 0.05 according to Mann Whitney U test.

### The effect of CS GoF mutations on glibenclamide sensitivity

Glibenclamide (glyburide) inhibits K_ATP_ channels in a biphasic manner, with high-affinity inhibition arising from interaction with the SUR subunit occurring at nanomolar to micromolar concentrations and low-affinity inhibition due to interaction with the Kir6.x subunit [32]. To specifically measure high affinity inhibition we applied glibenclamide up to 10 µM. Glibenclamide inhibited WT SUR2A K_ATP_ currents (in the absence of nucleotides), as well as D207E, Y981S, G985E and M1056I (Fig. 8), with maximal inhibition of ~70%, and IC_50_values ranging from ~15 to 45 nM. In contrast, mutations at residue 1150, in particular R1150W, resulted in a significant decrease in glibenclamide potency (Fig. 9).

**Figure 8:**
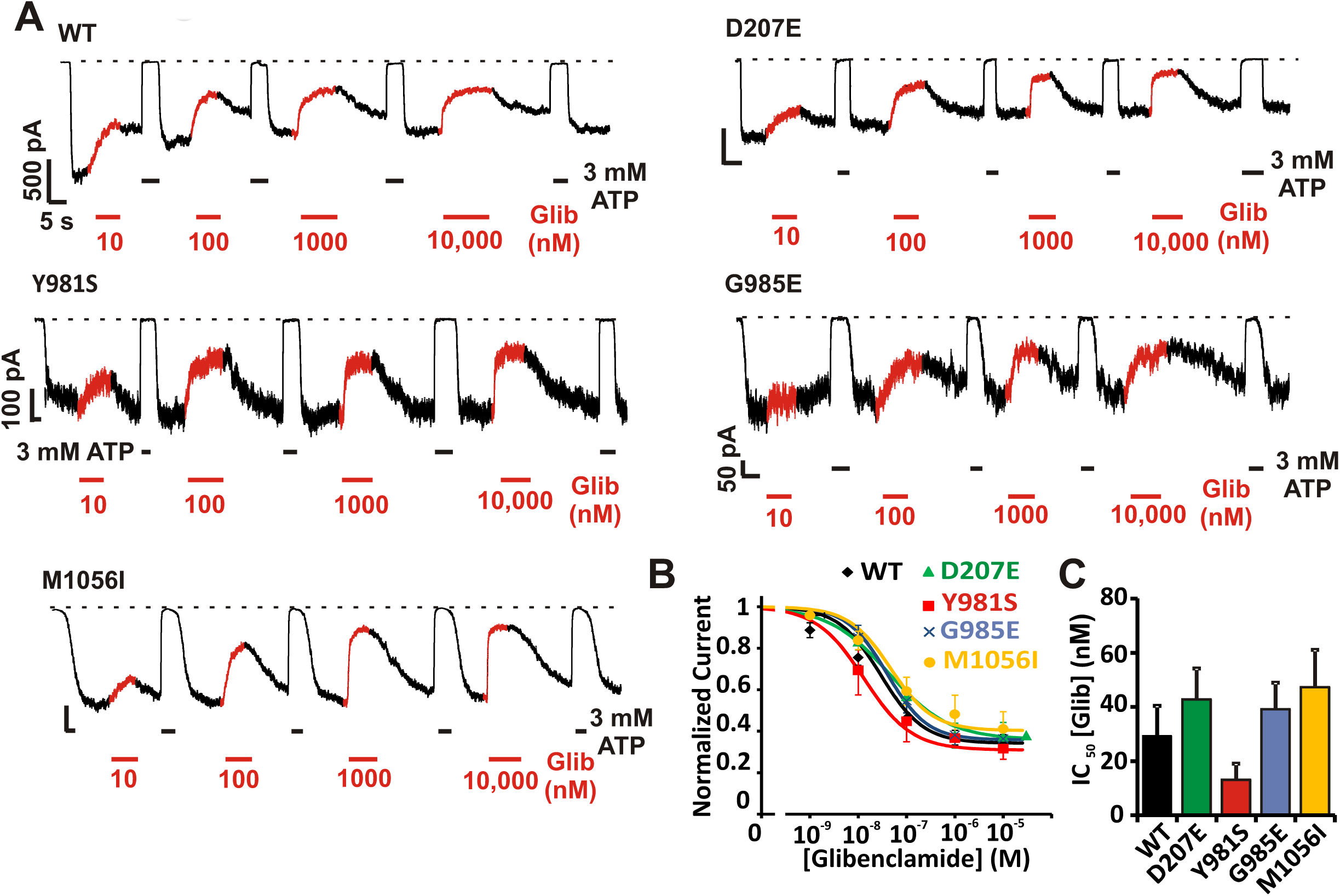
The D207E, Y981S, G985E and M1506I have no significant effect on glibenclamide sensitivity. Inside-out patch clamp recordings were made from Cosm6 cells transfected with Kir6.2 alongside WT or mutant SUR2A. Reversible inhibition was observed following administration of increasing glibenclamide concentrations, as shown in representative traces (scale bars denote 500 pA/5 s unless otherwise stated) **(A)**. Dose response analysis demonstrated that only minor, non-statistically significant variations in glibenclamide IC_50_ values were observed for these mutations (IC_50_ for SUR2A-WT 19.5 ± 6.1 nM, Hill coefficient 0.9 ± 0.1, n = 11; IC_50_ for D207E 42.8 ± 11.4 nM, Hill coefficient 0.7 ± 0.1, n = 4; IC_50_ for Y981S 13.1 ± 5.9 nM, Hill coefficient 0.9 ± 0.1, n = 3; IC_50_ for G985E 39.2 ± 9.8 nM, Hill coefficient 1.0 ± 0.2, n = 4; IC_50_ for M1056I 47.3 ± 13.8 nM, Hill coefficient 1.0 ± 0.4, n = 4;**) (B-C)**. Statistical significance was determined using Mann Whitney U tests.

**Figure 9:**
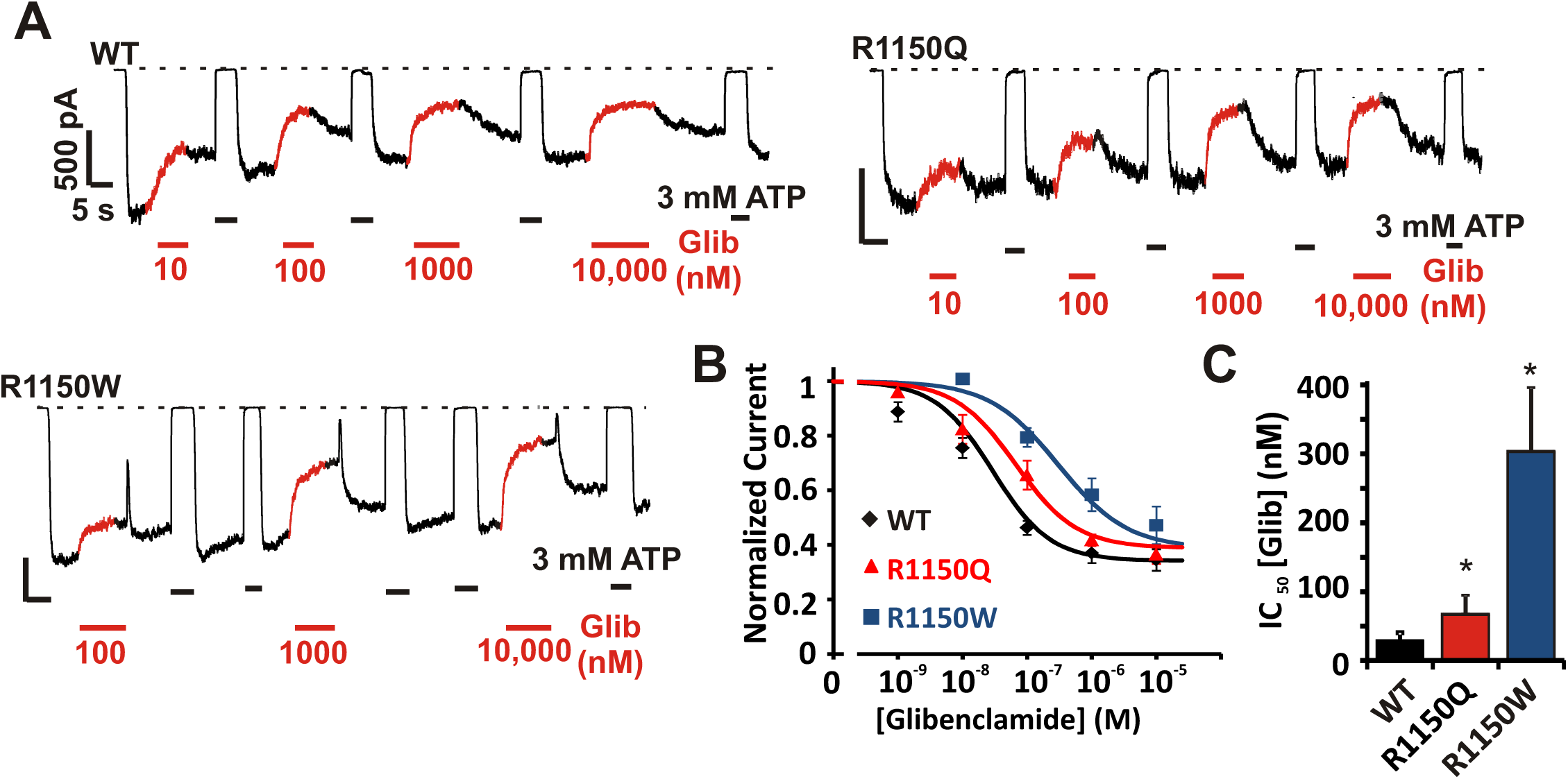
R1150Q and R1150W decrease glibenclamide sensitivity. Glibenclamide sensitivity measured from inside-out patch clamp recordings from Cosm6 cells expressing Kir6.2 with either SUR2A-WT, R1150Q or R1150W as shown in representative traces (scale bars denote 500 pA/5 s unless otherwise stated) **(A)**. Dose response analysis shows that both R1150Q and R1150W induce a statistically significant decrease in glibenclamide sensitivity (IC_50_ for R1150Q 86.3 ± 23.4 nM, Hill coefficient 0.8 ± 0.3, n = 5; IC_50_ for R1150W 303.5 ± 91.1 nM, Hill coefficient 0.8 ± 0.1, n = 5) **(B-C)**.

## Discussion

### Cantu Syndrome-associated ABCC9 mutations all cause K_ATP_ GoF

To date, the few analyzed Cantu Syndrome-associated mutations in *ABCC9* (SUR2), have been shown to result in gain-of-function of K_ATP_ channels in the presence of Mg^2+^ nucleotides, which can arise either from decreased sensitivity to inhibitory ATP, or augmented activation by Mg^2+^-nucleotides [16, 19]. In the present study we have demonstrated that four previously uncharacterized mutations in SUR2 (D207E [human D207E], Y981S [Y985S], G985E [G989E] and M1056I [M1060I]) lead to increased channel activity in the presence of regulatory nucleotides via diverse molecular mechanisms. As in previous reports, we used Kir6.2/SUR2A channels for analysis due to the difficulty of recording Kir6.1/SUR2 currents, although it is expected that the mechanism of SUR2 mutations will be conserved irrespective of the pore-forming subunit. In addition, although the dependence of channel activity on intracellular nucleotides differs quantitatively between the two major SUR2 splice variants (SUR2A and SUR2B, which differ only in their C-terminal exon) [33, 34], it is anticipated that, qualitatively, the changes observed for mutant SUR2A-containing channels will be common for SUR2B-containing channels. Interestingly, many CS features such as vascular dilatation and lymphedema likely arise from smooth muscle dysfunction, whilst effects on cardiac electrophysiology and skeletal muscle are less obvious [24]. This may suggest that the biophysical effects of mutations are more severe in SUR2B-than in SUR2A-containing channels. Alternatively, it is possible that the predominantly smooth muscle consequences of CS arise due to the physiological context of K_ATP_ function in smooth muscle rather than unique biophysical effects on SUR2B compared to SUR2A.

### The mechanistic basis of GoF varies between mutations

Here, we compared the sensitivity of wild type and mutant channels to ATP in the absence and presence of Mg^2+^, to separate Mg-independent inhibitory effect of ATP from the activating effect of MgATP. This analysis shows that the D207E mutation reduces ATP inhibition itself whilst the Y981S/G985E/M1056I mutations all increase activity by enhancing MgATP activation. The D207E mutation is found in the intracellular L0 linker between the two transmembrane domains TMD0 and TMD1 (Fig. 1). A critical role of the L0 linker in determining channel gating properties has been suggested for both SUR1 and SUR2 [35, 36]. Recent high resolution structures of the related Kir6.2/SUR1 K_ATP_ channel complex confirm earlier predictions that the L0 linker is closely apposed to the intracellular domains of the Kir6 subunit, and likely provides a structural interaction between the subunits that is involved in the transduction of functional signals. Interestingly, multiple mutations in the L0 linker of SUR1 have been reported in neonatal diabetes patients, including the analogous mutation to D207E, D209E [37]. Functional characterization of L0 linker mutations in SUR1 show that channel activity is increased due to elevations in the intrinsic channel open probability, arising from decreased occupation of long-lived, ATP-accessible closed states [35]. Consistent with this finding, we show that enhanced basal open probability of D207E-containing channels, reflected in decreased PIP_2_ activation, underlies decreased sensitivity to inhibitory ATP (Figs.4 & 5).

In contrast, we show that the Y981S, G985E and M1056I mutations all act by increasing K_ATP_ channel activation by MgATP (Fig. 6). These residues are all predicted to lie in close proximity to each other in a cluster within TMD2; Y981 and G985 are found at the N-terminal end of TM12, immediately following the NBD1-TMD2 linker, whilst M1056I is situated on the opposing TM13 (Fig. 1). The location of the Y981/G985/M1056 cluster at the link between the NBDs and the TMDs (Fig. 1) is appropriate for transduction of movements between the intracellular and transmembrane domains of SUR2. Biochemical analyses of SUR and related ABC proteins indicate that MgADP or MgATP binding to the NBDs of SUR may act to stabilize dimerization of NBD1 and NBD2 [7, 38, 39]. However, how binding or NBD dimerization is coupled to gating of the channel pore remains poorly understood.

In addition, we show that the previously reported common CS mutations R1150Q and R1150W (located in TM15 of TMD2) also enhance MgATP activation (Fig. 7), demonstrating that multiple transmembrane regions of TMD2 are involved in the conformational changes associated with Mg^2+^-nucleotide activation.

Notably, the gain-of-function induced by each mutation is quite subtle when mutant and WT SUR2A subunits are co-expressed to mimic the clinically relevant heterozygosity (Fig. 2). Recent reports of GoF mutations in Kir6.2 and SUR1 that underlie neonatal diabetes demonstrate that even subtle biophysical effects can result in disease [40], suggesting that dramatic changes may not be necessary. On the other hand, since SUR2Bs may be the more pathologically relevant splice variant, it is possible that these mutations will have a greater effect on channels containing SUR2B. In addition, the channel activity measured in 86Rb^+^ experiments under basal conditions may not fully recapitulate the metabolic and physiological context for K_ATP_ channels in muscle or other differentiated cell types, and so we cannot rule out a more significant activating effect under other conditions.

### Consequences for sulfonylurea sensitivity

Previous studies have demonstrated that second generation sulfonylureas such as glibenclamide inhibit SUR2-containing KATP channels, albeit with lower potency than SUR1-containing channels [41]. As such, glibenclamide, or other sulfonylureas, represents a potential pharmacotherapy for CS. However, there are multiple reports of neonatal diabetes mutations in the Kir6.2/SUR1 K_ATP_ subunits which reduce sulfonylurea sensitivity [28, 29], and as we have recently demonstrated, the CS-associated mutation V65M in Kir6.1 profoundly reduces glibenclamide inhibition of recombinant channels [30]. Therefore, it is important to assess the effect of SUR2 CS mutations on inhibitor sensitivity. Here, we show that the D207E, Y981S, G985E and M1056I mutations do not obviously affect glibenclamide sensitivity (Fig. 8), as determined in the absence of nucleotides in excised patch clamp experiments. It has been reported that sulfonylurea inhibition of SUR2-containing channels is affected by nucleotide regulation [42, 43], and so it is possible that these mutations may alter SU sensitivity under more complex physiological regulation, but this remains to be established.

A decrease in glibenclamide potency was observed in both the R1150Q and R1150W mutations (Fig. 9). Interestingly, R1150 lies in TM15 (Fig. 1) and previous studies have demonstrated that TMs14-16 are critical for high-affinity SU binding to SUR subunits [44, 45]. Indeed, serine to tyrosine substitution of a single residue in TM16 (predicted to lie within ~ 15 Å of R1150 on the cytoplasmic extensions of the TM helix) is sufficient to confer SUR2-like sensitivity to SUR1, and vice versa [44, 45]. This raises the possibility that the R1150 mutations may directly decrease glibenclamide sensitivity via disruption of the drug binding site. The R1150W mutation exhibited a more pronounced effect than the glutamine mutation at the same site, perhaps due to a greater steric effect of the bulky tryptophan sidechain. Regarding the relevance to treatability of disease, it is important to note that glibenclamide sensitivity was evaluated in a “homozygous” context where all SUR2 subunits were mutated, whilst CS patients identified so far are heterozygous for the gain of function mutations and therefore the effect of the R1150 mutations on SU sensitivity may be moderated in the patients.

Taken together, the present results provide further evidence for K_ATP_ gain of function consequences of SUR2 mutations in Cantu Syndrome. For several mutations clustered in TM12-13, the results illustrate a common mechanism (enhanced MgATP activation) without marked effect on sulfonylurea sensitivity. The results provide novel insights into the function of K_ATP_ channel complexes, can be useful for linking CS genotype to phenotype in this complex disorder, and will inform the consideration of therapeutic approaches to CS.

## Acknolwedgements

We thanks Ms. Risha Shah (Washington University) for help with Rb efflux experiments. This work was supported by NIH grant HL95010 to CGN. HIR is supported by the E-Rare Joint Transnational Cantu Treat program (I-2101-B26) to GvH.

